# Germinal centers are permissive to subdominant antibody responses

**DOI:** 10.1101/2023.05.31.543035

**Authors:** Philippe A. Robert, Theinmozhi Arulraj, Michael Meyer-Hermann

## Abstract

A protective humoral response to pathogens requires the development of high affinity antibodies in germinal centers (GC). The combination of antigens available during immunization has a strong impact on the strength and breadth of the antibody response. Antigens can display various levels of immunogenicity, and a hierarchy of immunodominance arises when the GC response to an antigen dampens the response to other antigens. Immunodominance is a challenge for the development of vaccines to mutating viruses, and for the development of broadly neutralizing antibodies. The extent by which antigens with different levels of immunogenicity compete for the induction of high affinity antibodies and therefore contribute to immunodominance is not known. Here, we perform *in silico* simulations of the GC response, using a structural representation of antigens with complex surface amino acid composition and topology. We generate antigens with different levels of immunogenicity and perform simulations with combinations of these antigens. We found that GC dynamics were driven by the most immunogenic antigen and immunodominance arose as affinity maturation to less immunogenic antigens was inhibited. However, this inhibition was moderate since the less immunogenic antigen exhibited a weak GC response in the absence of other antigens. Less immunogenic antigens reduced the dominance of GC responses to more immunogenic antigens, albeit at a later time point. The simulations suggest that increased vaccine valence may decrease immunodominance of the GC response to strongly immunogenic antigens and therefore, act as a potential strategy for the natural induction of broadly neutralizing antibodies in GC reactions.

## Introduction

The humoral immune response relies on the development of high affinity antibodies produced by plasma cells, and a reservoir of memory cells to potentiate the next encounters with pathogenic antigens. Affinity maturation happens in anatomical regions called germinal centers (GCs), where B cells can proliferate, mutate their B cell receptor (BCR) through somatic hypermutation (SHM), and go through mechanisms of selection, based on the affinities of their BCRs to available target antigens and their relative fitness compared to other B cells (Allen et al., 2007; Victora & Nussenzweig, 2022). After egress from the GC, B cells differentiate into plasma cells that secrete the soluble form of their BCR as antibodies, or become memory cells (Shlomchik & Weisel, 2012) that can persist in the body for several years.

The interplay between these complex mechanisms (Finney et al., 2018; Stebegg et al., 2018; Zhang et al., 2016) ensures that the host deploys antibodies to most pathogens. Antigens inducing such immune responses are termed “immunogenic”. Affinity maturation is sometimes suboptimal with viruses that mutate at a high rate. For instance, despite the endogenous production of high affinity antibodies to HIV, influenza virus or SARS-CoV-2, these antibodies are not necessarily neutralizing (Belongia et al., 2016; Haynes et al., 2022; Newman et al., 2022), and they may or may not have a sufficient cross-reactivity (i.e., the same antibody targeting multiple variants) or diversity (the mounted repertoire containing antibodies with diverse enough sequences to target different variants (Garg et al., 2023)) to cover the rapidly arising mutants. The evolutive forces between antibodies and antigens in play are complex, and predicting the outcome of a humoral response based on the combination of antigens and the prior history of infections is challenging but of major importance for the development of more efficient vaccination strategies.

The immune response to an antigen can be impacted by the simultaneous response to another antigen or another domain of the same antigen, which generates a hierarchy of immunodominance of GC responses to different antigens and their sub-domains. A canonical example is the antibody response to the influenza virus hemagglutinin (HA) protein, which contains highly accessible residues on its tip that induces a strong humoral response with high affinity antibodies. A functional binding domain forming a less accessible pocket mounts poor antibody responses, as well as a conserved stem domain with poor accessibility, and the remaining extracellular surface area covered by glycans that inhibit antibody binding (Doud et al., 2018; Wanzeck et al., 2011). In this context, the tip of HA induces “immunodominant” GC responses compared to the binding pocket which induces a suboptimal response and is called “subdominant”. The relative immunodominance between responses to HA domains is of therapeutic concern since the tip residues are not functionally important and are highly mutated providing a mechanism of escaping immunity. In contrast, the binding pocket is functional and tends to be conserved between variants. Therefore, antibodies targeting the binding pocket are more likely to have a large breadth to all variants since they structurally share this domain. However, conserved regions can still mutate (Chai et al., 2016), and there also exist broadly neutralizing antibodies targeting other, non-conserved domains of HA (Qiu et al., 2020). Apart from the antigen structure, the availability of BCRs (Abbott et al., 2018) that bind different domains of the antigen and their competition in the GC are also critical for the establishment of immunodominance. The availability of memory cells from previous immunizations (Sokal et al., 2023) can mitigate the relative response to different antigens. It has also been proposed that the memory response to a previous antigen variant might decrease the strength of recognition to newer strains by skewed recognition to those antigens, termed the “Original Antigenic Sin” (Francis, 1960) while computer simulations suggested that memory responses would promote a shift to antibody responses to less immunogenic epitopes (Meyer-Hermann, 2019).

The evolution of B cell clones in GCs often leads to the clonal dominance of only a few B cell clones (Garg et al., 2022; Meyer-Hermann, 2021; Tas et al., 2016), which bind epitopes in more immunogenic domains. Interestingly, low affinity B cells have been observed to participate and persist in GCs despite the presence of high affinity cells (Kuraoka et al., 2016), which suggests that GC responses to weakly immunogenic epitopes can evolve despite the presence of an immunodominant concomitant response. This raises two questions: i) Is a GC response to a particular antigenic domain subdominant due to the presence of another highly immunogenic antigenic domain, or due to absolute poor immunogenicity? In the former case, immunodominance is relative, and shielding targeted epitopes driving the immunodominant response by blocking antibodies could rescue the GC response to the other antigenic domain, as suggested in (Meyer-Hermann, 2019). ii) How to modulate the composition of antigens in vaccines to potentiate the response to antigenic domains raising subdominant responses?

Interestingly, repeated immunizations with identical (Saunders et al., 2017) or mutated (Escolano et al., 2016; Tian et al., 2016) antigens enhance the development of broadly neutralizing antibodies, suggesting that the permissive selection of low affinity B cells in GCs can be used to target less immunogenic epitopes by repeated immunization. The reasons for successful amplification of responses to such less immunogenic epitopes are not known. A combination of a large set of representative peptides covering influenza HA strains over a hundred years was capable of raising antibodies with high breadth, suggesting that the dilution of highly immunogenic antigens by antigen cocktails may overcome the barrier imposed by immunodominance (Carter et al., 2016). The low frequency of naive B cells capable of binding to less immunogenic epitopes is also considered as a factor limiting the induction of broadly neutralizing antibodies (Abbott et al., 2018), in which case mutating the antigen to better catch the rare B cells was a proposed strategy (Escolano et al., 2016).

Additional evolutionary forces at longer time scales and between multiple GCs have also been suggested to impact on immunodominance, such as the antibody feedback (Zhang et al., 2013) where previously accumulated antibodies may shield certain antigen epitopes in the GC (Meyer-Hermann, 2019), or the co-evolution between B-cell lineages and virus *in vivo (Gao et al., 2014; Liao et al., 2013)*. For instance, in the context of malaria vaccine, existing serum antibodies inhibited the reactivation of memory B cells after the third vaccine shot (McNamara et al., 2020). This ongoing discussion emphasizes that a quantitative understanding of the interplay of forces responsible for immunodominance is needed and of therapeutic importance.

Mathematical modelling has extensively been used to study the dynamical properties of the GC, to test potential mechanisms behind experimental observations that were later validated, and to propose targeted modulations of affinity maturation (Buchauer & Wardemann, 2019). The properties of competition between antibodies targeting overlapping epitopes (Yan & Wang, 2020) and co-evolution with virus mutations (Sheng & Wang, 2021) have also been modelled. We showed *in silico* that combining multiple antigens that differ by only a few mutations increased the cross-reactivity of produced antibodies (Robert et al., 2021). In vaccine strategies that use mutated HIV variants containing antigenic regions that lead to both an immunodominant and subdominant response, it was suggested *in silico* (Wang, 2017; Wang et al., 2015) that a higher antibody cross-reactivity is achieved using sequential rather than cocktails immunizations. This was shown in a simulation setting where B cells can only detect one of the antigens at the selection step, i.e. where antigen dilution mechanistically decreases chances of B cell survival. Simulation of the GC response to multiple HA influenza variants suggested that the choice of antigens for the first immunization can reduce the diversity of available BCRs and decrease the chance of broad neutralization in the next immunizations (Amitai et al., 2020). Increasing the number of unrelated antigens (valency) during immunization (Childs et al., 2015) was predicted to allow for high affinity GC B cells, at the expense of reduced number of plasma cells recognizing each antigen compared to lower valency immunizations. Low immunodominance of responses to rare antigens was rescued *in silico* by the injection antibodies against the epitope dominating the GC response (Meyer-Hermann, 2019).

Note that the aforementioned studies were based on various abstract settings (Robert et al., 2018), which limits their predictive power. Here, we aim at investigating immunodominance in a more realistic setup by using a structural representation of synthetic 3D antigens, which naturally enables the development of antibodies against different epitopes and with different strengths depending on their surface amino acid composition and topology. We asked under which conditions immunodominant responses emerge when two or more antigens are present, and specifically, how the level of immunogenicity of an antigen contributes to immunodominance in the presence of other antigens. We selected different classes of antigens with different levels of repertoire recognition as the basis for different classes of immunogenicity, and asked to which extent immunodominance arises from intrinsic antigen immunogenicity or by the combination of antigens used.

We observe that the presence of a more immunogenic antigen only moderately inhibits the response to a less immunogenic antigen in the same GC allowing a substantial response to the latter. Further, in later days of the GC development, lower immunogenic antigens dampen affinity maturation to more immunogenic antigens present in the GC. This suggests that immunodominance of GC responses to different antigens only modulates but does not abrogate the immune response to antigens with low immunogenicity that would intrinsically induce a poor response. These results suggest that an increased antigen valency in vaccines would contribute to dampen the response to antigens dominating the GC response.

## Results

### Generation of an antigen library with different GC response strength

In order to simulate GC reactions where different degrees of immunodominance naturally emerge, we generated antigens of different immunogenicity. We used an agent-based model of the GC, that simulates migration, proliferation, mutation, cell-cell contacts and selection (Meyer-Hermann et al., 2012; Robert et al., 2021) (Figure 1A). The mechanistic B cell decisions rely on the BCR affinity to the available antigens, which require a formula to estimate the affinity between a BCR sequence to every antigen. Since antibody-antigen affinity prediction is not feasible at the scale of a GC simulation, which necessitates the assessment of affinities of 10,000 to 100,000 mutated BCRs to each antigen (Robert et al., 2021), all GC models have used simplified affinity models, ranging from the shape space, to binary sequences, cubic proteins, and lattice 3D protein structures (Robert et al., 2018). In the simpler models, immunodominance was manually implanted, for instance by giving a penalty to antibodies binding a pocket-like conserved domain (Wang et al., 2015) or by fixing antigens of different amounts at specific positions in the shape space (Meyer-Hermann, 2019). These methods were suited to study the downstream competition between clones and ultimately predict conditions that maximize the generation of antibodies against a predefined domain. Here, we use a more realistic structural model where BCR affinity is defined by exploring all possible 3D conformations of the BCR binding domain (Complementarity Determining Region 3 from the heavy chain, CDRH3) to a 3D antigen structure. Antigens and antibodies were modelled on a 3D lattice, and affinities of BCRs were calculated as the fitness of the energetically optimal BCR structure to the antigen structure (Figure 1B). Instead of manually designing antigen structures with hidden or accessible surfaces as done in previous studies, we took the structural topology of real-world proteins as a source of 3D structures. For that purpose, structures from a Protein Data Bank (PDB) were discretized and projected on the 3D lattice. This formalism allows for complex properties to be recapitulated without the need to be manually imposed. Sequences representing CDRH3 are randomly sampled to generate founder B cells, and whenever a mutation occurs, the affinity of the BCR towards all antigens considered in the simulation is re-evaluated (see Methods).

**Figure 1:**
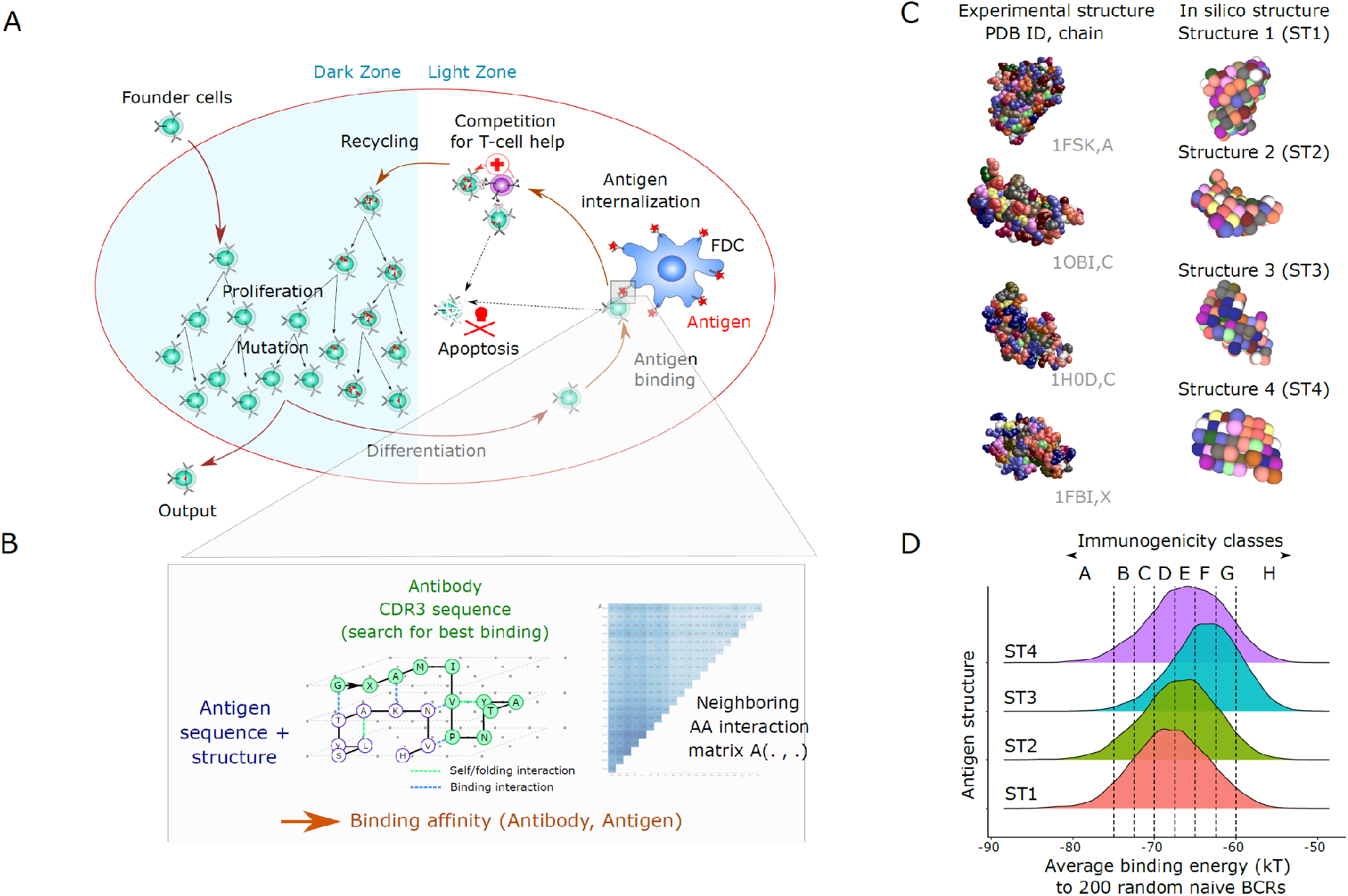
Mathematical model of the GC response and design of complex antigens with different levels of immunogenicity. **A**. Biological mechanisms included in the 3D cellular model: founder cells with random BCR sequences proliferate and mutate by SHM in the dark zone, migrate to the light zone where they compete for binding antigen displayed on Follicular Dendritic Cells (FDCs). Depending on the BCR affinity to the antigens present in the GC, the B cell can capture antigen, process it and present it to T follicular helper cells (Tfh). B cells that receive sufficient signals from Tfh cells recycle to the dark zone and initiate additional rounds of proliferation and mutation. **B**. An antigen is represented as a 3D structure on a lattice with one amino-acid per node (Robert et al. 2021). The BCR sequence is folded according to every possible binding conformation around the antigen structure (exhaustive docking). The binding energy of each possible BCR-antigen binding conformation is defined by an empirical energy potential for each neighbor pair of amino acids between the BCR and the antigen. The energetically optimal structure defines the binding energy (in kT units), further converted into a unitless binding affinity. **C**. Design of antigens. We used a discretization method (Latfit, see methods) to convert the experimental structure of four different proteins from the PDB into *in silico* antigen structures ST1-4. Then, we designed on those antigen structures many antigens that vary by their amino acid surface composition. **D**. Antigen immunogenicity classes are defined by the strength of recognition by naive B cells (x axis). The graph shows the average recognition (binding energy) of 6000 antigens with random amino acid composition by 200 random naive B cells for antigen structures ST1-4. The distribution reflects the effect of changing the surface amino acid composition of antigens. PDB: Protein Data Bank, BCR: B-cell receptor, GC: germinal center.

Amino acid modifications of the vaccine antigen can enhance the recognition by naïve B cells (or a subset thereof) (Escolano et al., 2016). In our simulation, we first investigated to what extent the surface amino acid composition impacts the GC response. Specifically, we used four antigen structures, named ST1 to ST4, discretized from PDBs 1FSK, 1OB1, 1H0D and 1FBI, chains A, C, C and X, respectively (Figure 1C), and randomly assigned residues to the surface of these structures to generate many antigens with complex 3D structure and different amino acid composition (Figure 1C).

We randomly generated 200 sequences to represent naive BCR sequences and assessed their average binding energies with 6000 modified antigens per antigen structure (Figure 1D). The 200 sequences were a good representative of a large pool of 10,000 BCR sequences in predicting the average binding energy (Supplementary Table 1). The average binding energy ranged from -85 to -55 kT depending on the amino acid composition of the antigen. This reflects that the amino acid composition of antigens alone could alter the recognition by naive B cells, i.e., its immunogenicity. The scaffolding structure also had an impact, as antigens with structure ST3 had higher average binding energies (i.e., lower affinity) while antigens with structure ST1 had lower average binding energy, showing that our simulation framework accounts for the contribution of structural properties to immunogenicity. We assigned *in silico* classes of immunogenicity to the antigens based on their average affinity to BCRs (Figure 1D), ranging from the high affinity A class (average energy less than -75 kT units) by steps of 2.5 kT up to the remaining low affinity H class (average energy higher than -60 kT units).

In order to see how antigens from the different immunogenicity classes A-F translate into GC responses, we performed GC simulations with randomly selected antigens from classes A-F using structure ST3, named A3-F3 (Figure 2), and the other structures (Supplementary Figure 1). Antigens A3 to D3 enabled a strong GC response, as observed by a higher GC volume (Figure 2A), higher affinity of GC B (Figure 2B) and produced output cells (Figure 2C). In comparison, antigens with low immunogenicity (E3 and F3) succeeded in inducing a GC despite a very low affinity maturation of GC B or output cells (Figure 2A-C), showing that a response to most available antigens across a large range of immunogenicity was mounted in the simulations.

**Figure 2:**
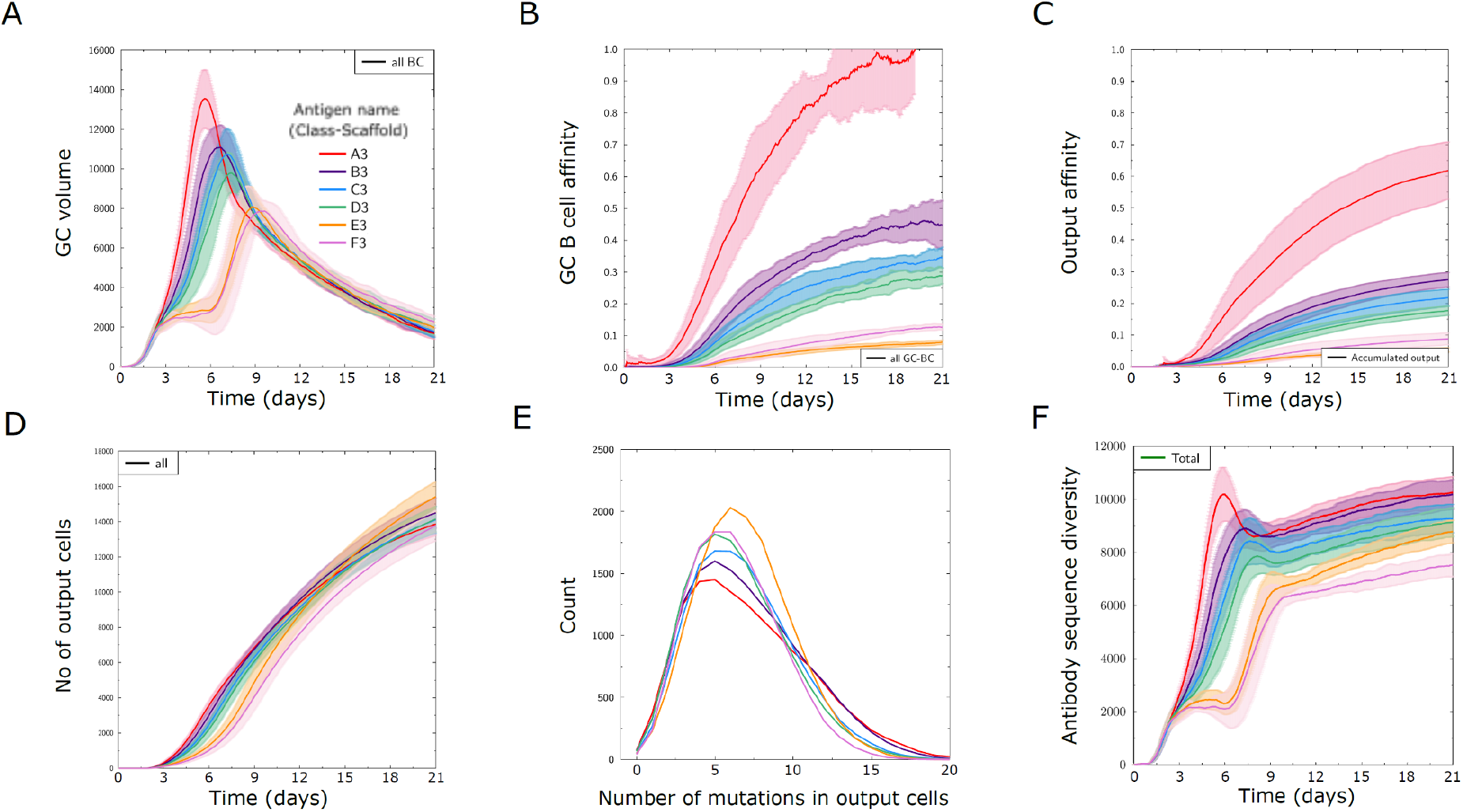
The designed classes of antigen immunogenicity mirror different strengths of GC responses. Results of GC simulations with an antigen from each immunogenicity class with the scaffold structure ST3 (A3 – F3) as defined in Figure 1D. **A**. Volume of GC measured as number of GC B cells over time. **B**. Average affinity of B cells in the GC. **C**. Average affinity of output cells. **D**. Number of output cells produced by the GC. **E**. Integral distribution of number of mutations (SHM) in output cells in all simulations. **F**. Antibody sequence diversity, measured as the number of distinct antibody sequences present in the GC over time. Solid lines and shaded areas represent the mean and standard deviation of 10 independent GC simulations, respectively.

Interestingly, there were only minor differences in the number of output cells produced using antigens of different classes (Figure 2D). Output cells shared a similar number of mutations with a wider distribution for antigens with high immunogenicity (Figure 2E), and the diversity of mounted BCR sequences was lower for antigen classes with less immunogenicity (Figure 2F). This shows that the antigens designed with different immunogenicity levels can be employed to study conditions of relative immunodominance between antigens: The reasons for differential GC response of antigens from classes A to F are multifactorial, ranging from the properties of naive B cells to the fact that some antigens are harder to bind for structural reasons, thus creating different mutational landscapes. The simulations also allow to follow the antigen domains bound by BCR sequences over time (Figure 3).

**Figure 3:**
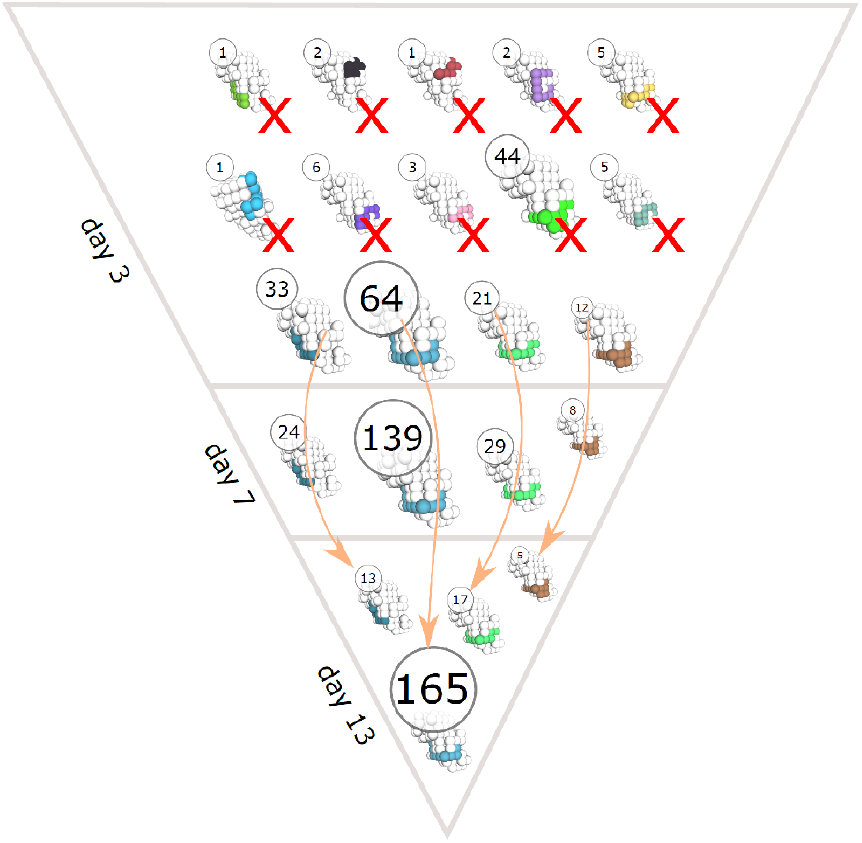
Time evolution of clonal dominance and antigen domain recognition of BCRs in a representative GC simulation. The structure of the antigen is shown in white and the binding conformations of BCR sequences are colored. The circles show the number of unique BCR sequences that bind the antigen with the corresponding binding conformation. Arrows indicate changes in the abundance of BCR sequences with the same binding conformation at different time points (day 3, 7 and 13 of the GC simulation) and cross marks (X) indicate binding conformations that got extinct during the GC reaction.

### Observed dynamics of GC responses are defined by the antigen of highest immunogenicity

We followed dynamics of GCs containing two antigens of different immunogenicity (B3+D3 or D3+F3), taking “B” as high, “D” as medium and “F” as low immunogenicity classes (Figure 4 for structure ST3). Simulations with two antigens started with 50% of randomly selected BCR sequences for GC founder cells that recognize each antigen with a minimum affinity. Each antigen was represented in equal amounts on the FDCs. Therefore no antigen is favored by naive cell frequencies or antigen amount by design.

**Figure 4:**
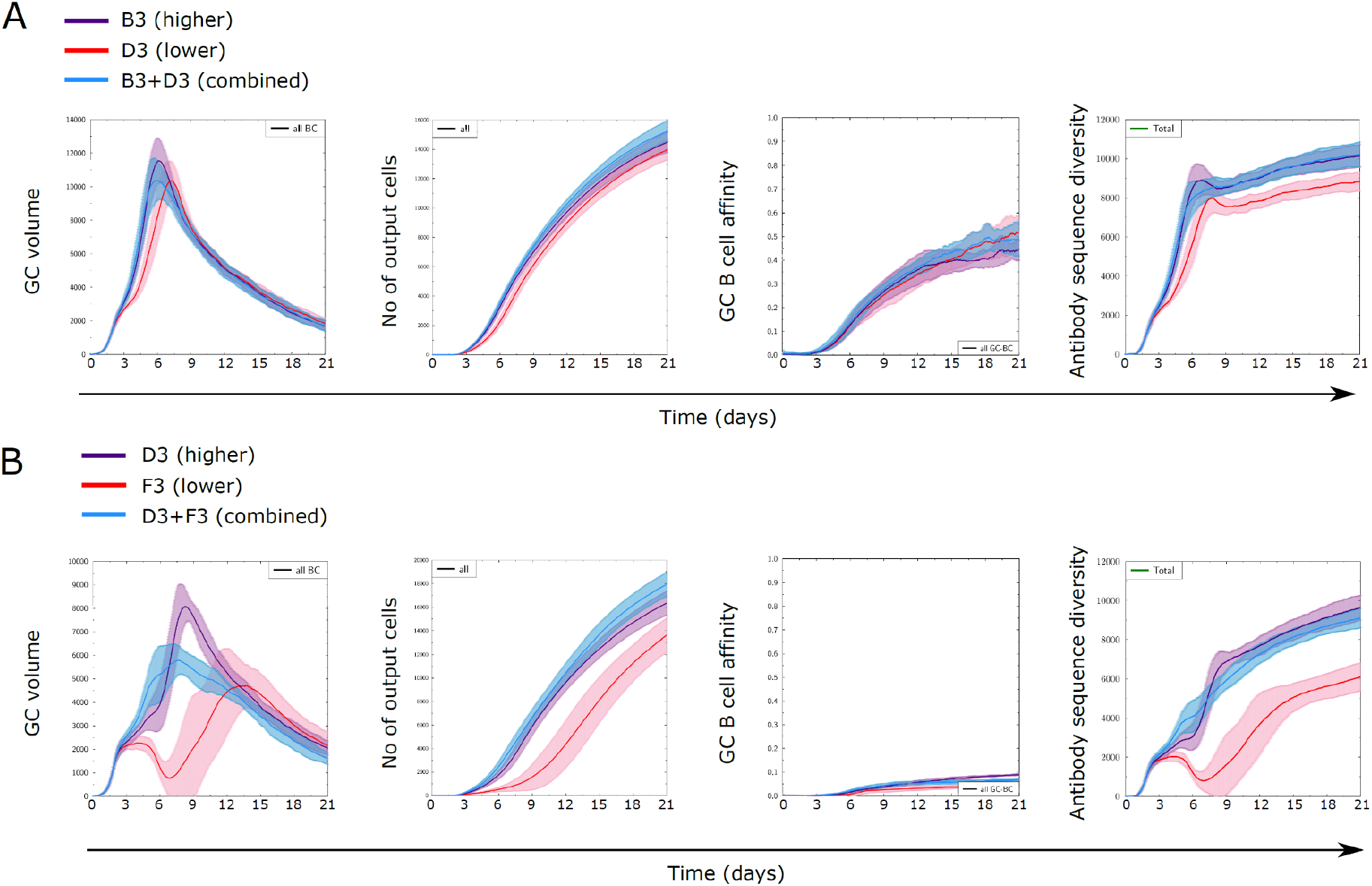
GC dynamics with two antigens of different immunogenicities in equal amounts. Control simulations with only one antigen with higher (purple) or lower (red) immunogenicity and with both antigens combined (blue). **A**. GC simulations with antigens of immunogenicity B and D versus **B**. for immunogenicity D and F. Because simulations are performed with the same initial amount of antigen, simulations with two antigens are started with half the dose of each antigen.

Simulations combining B and D antigens (B3+D3) showed GC dynamics similar to simulations with B alone, in terms of GC volume, affinity and diversity, but differed substantially with D alone (Figure 4A). Similarly, simulations combining D and F (D3+F3) antigens were similar to simulations with D antigen alone but for a slight reduction in the GC volume peak with both antigens, but different from simulations with F alone (Figure 4B). This shows that for two antigens the observed GC dynamics was mainly driven by the most immunogenic antigen present within a GC. These findings were consistently found for other antigen structures (Supplementary Figure 2) and suggest that observing the GC dynamics alone does not suffice to inform about the antigenic composition present in a GC nor about the development of subdominant responses.

### More immunogenic antigens permissively inhibit the response to less immunogenic antigens

Since the GC dynamics was driven by the most immunogenic antigen, we asked to what extent the GC was permissive to the development of a GC response to the other antigen.

We considered the simulation with two antigens D and F to analyze the distribution of binding affinities of GC B cells to both antigens at the late phase of affinity maturation (day 13), for antigens on each structure (Figure 5). As a control, we used simulations with a single antigen to identify the range of affinities in GCs unperturbed by a second antigen. We could identify populations with high affinity to D and F in the respective single antigen simulations. Depending on the structure, multiple populations coexisted with different ranges of affinities to B and/or D, and in the case of structure 3, simulations with F raised antibodies with medium-high affinities to D as well, suggesting that for some antigen combinations, a potential sharing of epitope features makes it possible for antibodies generated by a single antigen to recognize another antigen. Interestingly, simulations with D and F showed the presence of two or more populations, containing populations skewed to recognize D and populations skewed to recognize F. This means that the GC simulation was permissive to the development of B cells specific for both the high and low immunogenic antigen.

**Figure 5:**
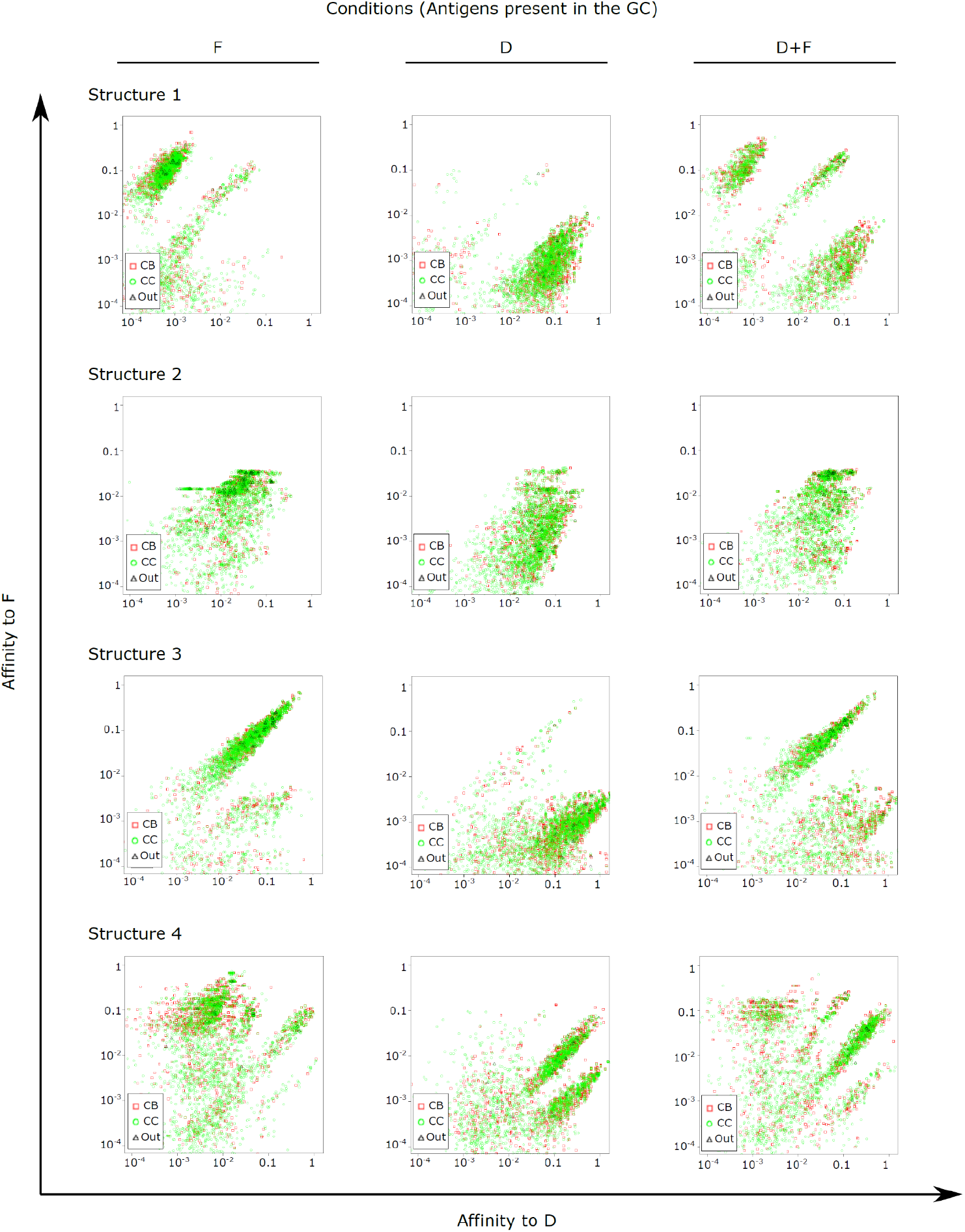
Distribution of affinities of single GC B cells to antigens of classes B and D at day 13 of the GC simulation. Simulations were performed in the presence of either or both antigens as indicated above the figure panels. Points represent single B cells. One representative simulation out of 10 independent simulations is shown for each antigen structure.

We then quantified the average affinity of B cells towards antigens of immunogenicity class D as a function of the presence of other antigens, either of class B (higher) or F (lower immunogenicity), to understand whether the presence of more or less immunogenic antigens can inhibit the development of GC response to the antigen of interest (Figure 6A). We also displayed the affinities to D relative to affinities at the beginning of simulation (Figure 6B), to account for the differences in their affinity dynamics between antigens on different structures.

**Figure 6:**
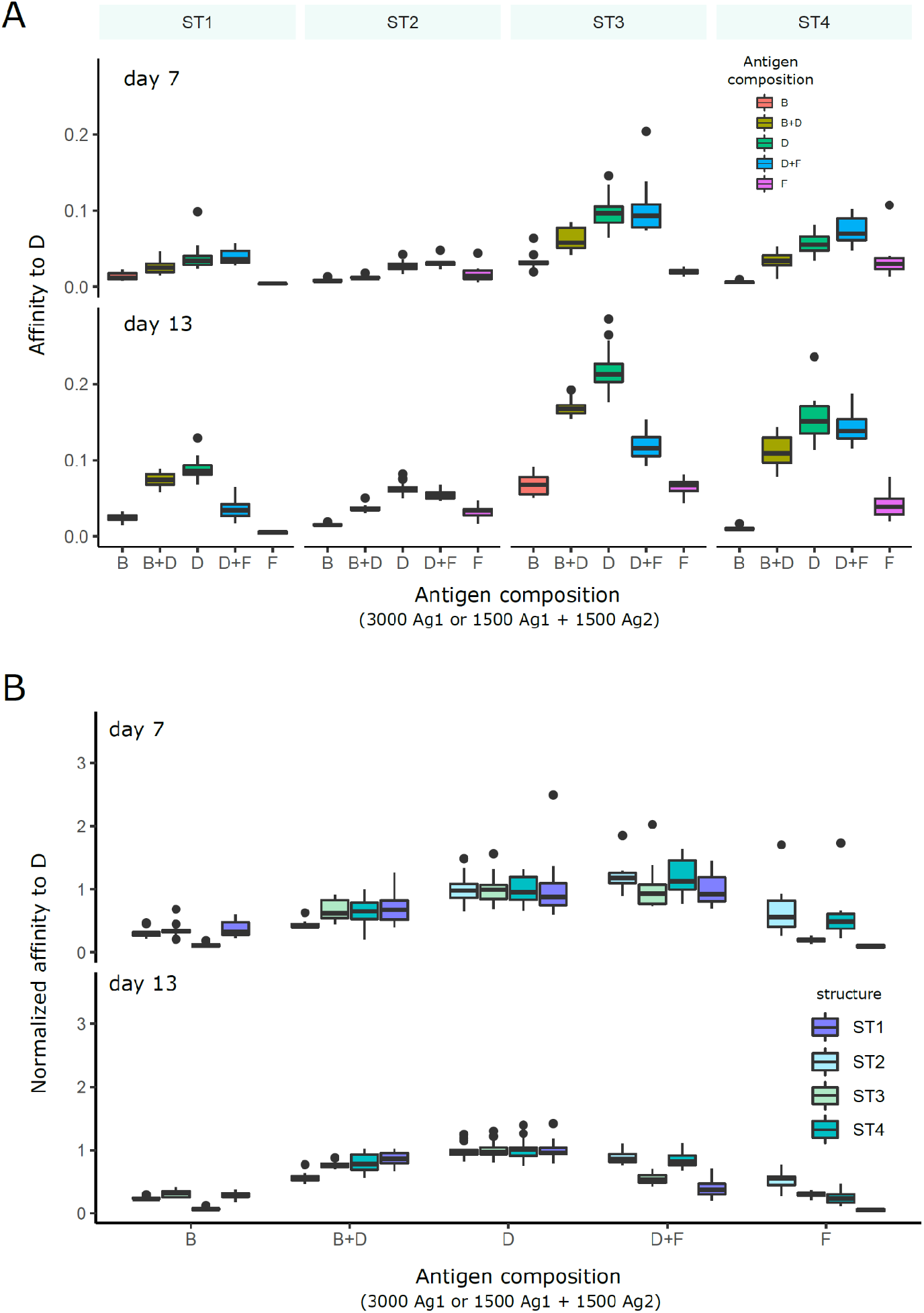
Affinity of GC B cells to antigen D in GC simulations with different combinations of antigens presented in the GC. Each condition was performed 10 times and the box plots show the median and 25% quantiles. **A**. average affinity of GC B cells to D for simulations with each antigen structure separately. **B**. affinities normalized to the average affinity to D in the simulation with D alone in order to account for the effect of each antigen structure on the GC responses. Normalization was done for each structure separately. The different antigen structures ST1-4 are color coded (legend). In simulations with antigen B or F alone, D is absent and no strong affinity maturation to D is expected. In conditions with two antigens (B+D and D+F), the total amount of initial antigen is equally distributed on each antigen.

Using simulations with only B or F antigen as reference, affinity to D stayed low, which is expected since D was absent, and D, B and F have unrelated sequences. We then added antigen of class D to the simulation. At an early time-point (day 7), affinities to D were low in the presence of B, but not in the presence of antigen of class F with lower immunogenicity. However, at a later time-point (day 13), affinity to D was decreased both in the presence of B or F. This suggests that immunodominance develops over time: Antigens with a certain level of immunogenicity inhibit the development of GC response to less immunogenic antigens, but not to more immunogenic antigens in the initial phase. Later, less immunogenic antigens may “catch-up”. This also means that an immunodominant response to an antigen is permissive to the concomitant response to other antigens, and suggests that antigens with low immunogenicity can dampen the GC response to highly immunogenic antigens at later time-points.

### Antigens of low immunogenicity dampen affinity maturation to more immunogenic antigens independently of antigen dilution

To test whether the reduction of antigen dose in simulations with two antigens was responsible for the inhibition between concomitant GC responses, we performed simulations with single antigens at half-dose (Figure 7) and compared them to simulations with either single antigen at full dose or two antigens with half dose. Interestingly, affinity to D was increased with half the amount of antigen of class D, which reflects increased competition for antigen. This implies that the reduction of affinities in the presence of other antigens is not due to a reduced amount of antigen.

**Figure 7:**
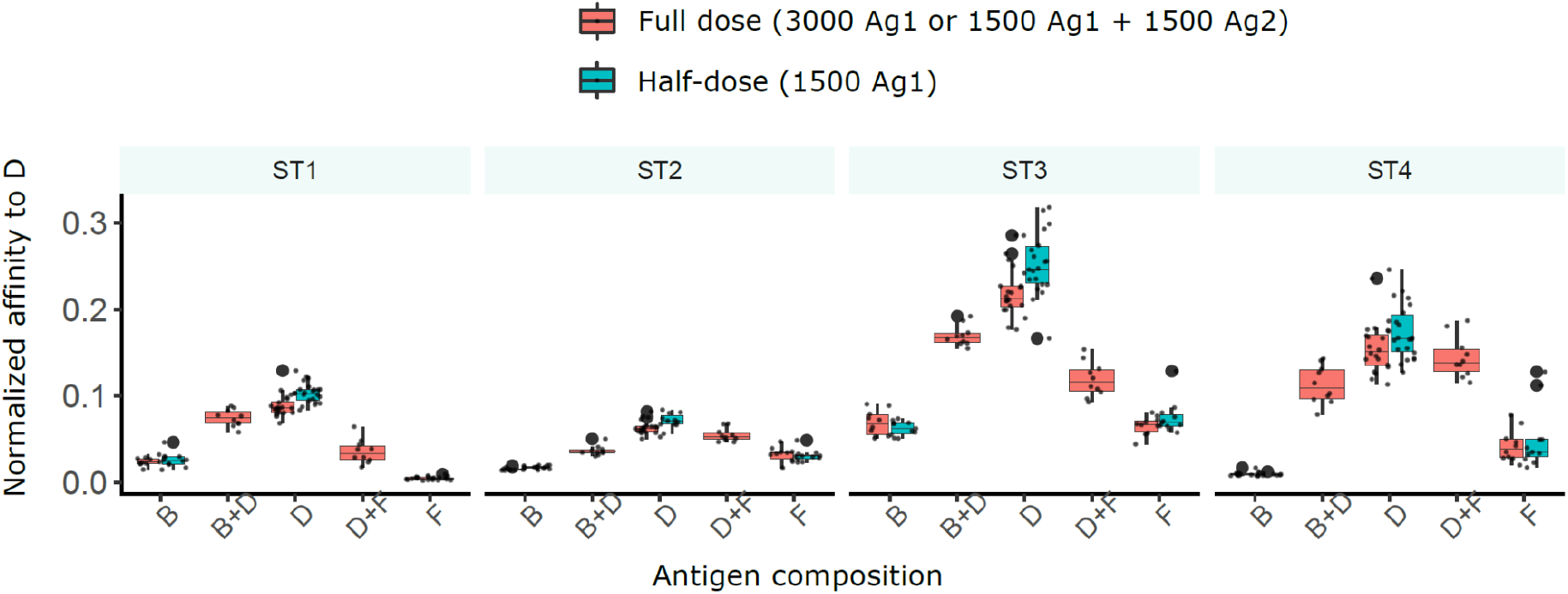
Antigen amount is not the reason for reduced affinity to D in the presence of other antigens. Simulations with two antigens (B+D or D+F) and 1500 units of each antigen (full dose, red), or with one antigen (B or D or F) and 3000 units of antigen (full dose, red) or 1500 units (half-dose, turquoise). 10 simulations are shown per condition at day 13, the barplots show median and 25% quantiles.

### Impact of antigen valency of immunizations with antigens of different immunogenicity

Since the presence of a highly immunogenic antigen did not completely abrogate the GC response to less immunogenic antigens, we investigated to which extent one could combine multiple antigens in order to decrease the dominant response to the more immunogenic antigen, and how many different such antigens a GC response could bear without losing the capacity to produce antibodies to all antigens. We performed simulations where one class D antigen was combined with 1 or more class F antigens with equal amounts and specific founder cells for each antigen (Figure 8). In terms of GC dynamics, the addition of multiple low immunogenic antigens decreased the strength of the GC response while slightly increasing the number of output cells, suggesting that the dominant GC response to D was reduced with increasing antigen valency.

**Figure 8:**
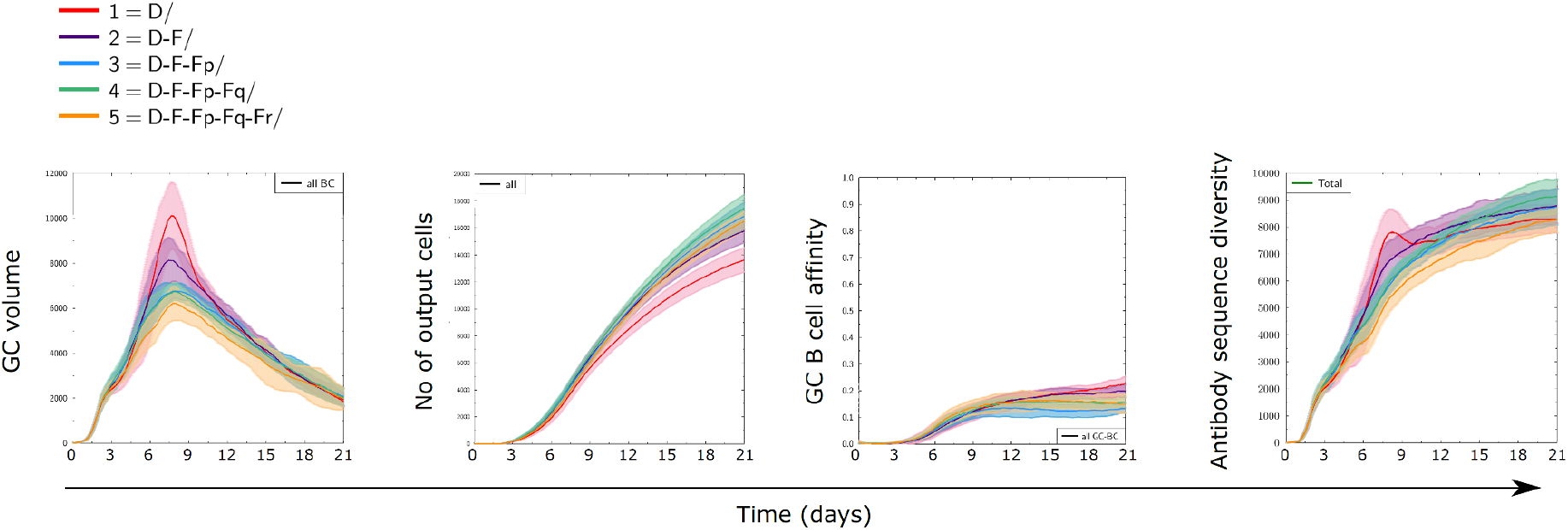
Dynamics of GC responses to combinations of antigen of class D with one or more antigens of class F. In addition to antigens D (D3) and F (F3), antigens Fp, Fq, and Fr of equal recognition by naive B cells were randomly generated. There was no sequence similarity between any of those (Supplementary Table 2). For instance, the combination D+F+Fp+Fq contains four antigens, whose initial amounts were divided equally to reach a total of 3000 units at start. Equal fractions of founder cells were selected to recognize each antigen at start. All antigens were defined on structure ST3. Solid lines and shaded areas represent mean and standard deviation of 10 GC simulations.

## Discussion

Although immunodominance is considered as a challenge for vaccine development, the relative effect between the intrinsic immunogenicity of antigens and relative immunodominance of the GC response is underexplored. We observed that the amino acid composition of antigens and their structural properties can modulate the strength of the GC response. We designed a strategy to generate antigens with different levels of immunogenicity based on their average affinity to naive B cells and used it as a model system to study immunodominance in GC responses. In particular, this model allows us to quantitatively compare the strength of mutual inhibition between GC responses to multiple antigens, and to investigate reasons or mitigations for this process. In immunizations with cocktails of antigens with different immunogenicity, we observe that the GC response to each antigen is mainly driven by the immunogenicity of the antigen, although competitive effects also play a role. The GC dynamics were mainly driven by the antigen of highest immunogenicity but remained permissive to antigens with lower immunogenicity, showing that an observed strong GC response does not exclude the parallel response of the GCs to low immunogenicity antigens (Kuraoka et al., 2016; Silver et al., 2018). We provided evidence that low immunogenic antigens can not only survive but are capable of partially inhibiting the GC response to more immunogenic antigens at late time-points.

Immunogenicity is generally defined as the capacity of the antigen to mount an immune response (Ilinskaya & Dobrovolskaia, 2016) and is a combined property of both the antigen and the immune system irrespective of the presence of other antigens. In different contexts, immunogenicity may refer to different stages of the immune response. Here, we have defined immunogenicity directly at the level of naive repertoire recognition, which is the earliest level of specific recognition by the adaptive immune system and has already been shown to be critical for GC responses (Abbott et al., 2018). We have also shown that the strength of the GC response directly correlates with the degree of repertoire recognition, justifying the definition of “immunogenicity classes based on the level of recognition by naive B cells (Figure 1). However, antigens may exist that are well recognized by naive B cells while not mounting a strong GC response, for instance by lack of specific T cells, or if antigens are similar to self-antigens and the GC response is inhibited by complex mechanisms such as clonal redemption (Burnett et al., 2018).

The term immunodominance is often ambiguously used without explicitly mentioning antigen combination nor pre-existing immune state. Further, antigens are sometimes called subdominant or immunodominant, which is ambiguous since the very same antigen might raise subdominant or immunodominant responses depending on the presence of other antigens. Therefore, we distinguished the antigen-intrinsic property of immunogenicity from immunodominance, which qualifies the GC response for a specified antigenic setting. Here, we considered the general case of immunodominant-subdominant responses between two or more domains of the same antigen during a primary GC response. Of note, immunodominance can also observed at a smaller scale between responses to different epitopes of the same domain, for instance due to steric hindrance between antibodies binding physically close epitopes (Yan & Wang, 2020), or between the antigen of interest and bystander antigens present in the GC such as self-antigens or commensal antigens in Peyer’s patches, which relates to the general case of multiple unrelated antigens. While the structural BCR-antigen binding framework allows for the emergence of a BCR with cross-reactivity to unrelated antigens (Robert et al., 2022), in most cases cross-reactivity will not be a major driving factor of GC responses to multiple unrelated antigens.

Multiple factors contribute to immunogenicity of antigens such as the nature of antigen, antigen dosage, the number and affinity of antigen specific naive B cells. In this study, we modulated the antigen amino acid composition in order to achieve different repertoire recognition and to define immunogenicity classes. To eliminate the effect of other factors, we performed simulations in controlled settings by considering equal amounts of each antigen and similar numbers of antigen specific founder B cells with affinities above a fixed threshold. However, antigens with low immunogenicity may activate less naive B cells, and therefore the amount and affinity of founder cells might also contribute to immunogenicity of antigens, and could be further simulated as an additional factor. In the simulations, the mutation landscape is different between the classes of antigens, which infers differences in the rate and extent of B cell affinity maturation to a particular domain. Mutations of B cells to an immunogenic antigen may reach higher affinities faster (since the mutation frequency depends on affinity (Gitlin et al., 2015)), while mutations to a low immunogenicity antigen might be bounded to lower affinities, or require more mutations to reach high affinities in comparison to the former antigen. This highlights the advantage of using a structural model for affinities, since many such properties are inherently reproduced eliminating the need for manual incorporation.

In the GC model, we assume that Tfh cells can provide survival signals to any B cell irrespective of the antigen they recognize, which is relevant when antigens represent subdomains of the same protein, since when B cells internalize one domain, they also internalize the other domains of the same antigen and present the same set of peptides of the full antigen to Tfhs. Inclusion of Tfh cells with different specificities will enable assessing the impact of Tfh repertoire in the establishment of immunodominance, with potential clonal competition between Tfhs since Tfh survival is also conditioned by T-B interactions (Baumjohann et al., 2013), which goes beyond the general scope of this study.

In this study, we focus on investigating the immune response in primary GC responses and therefore neglect the presence of memory B cells, pre-existing serum antibodies and the contribution of an altered immune properties such as age (Linterman et al., 2023) or the degree of chronic inflammation. Although it is clear that memory cells participate in secondary GC reactions (Sokal et al., 2023), the extent of participation is unclear (Mesin et al., 2020). In our previous study (Robert et al., 2021), considering 5% of founder B cells as memory B cells substantially increased the affinity of output cells. Future studies investigating secondary GC reactions by injecting different fractions of memory B cells as founder cells would help understand whether memory B cells amplify or inhibit the subdominant responses, depending on their antigen specificities. The process of antibody feedback (Zhang et al., 2013) where pre-existing serum antibodies compete with B cells for antigen binding in the GC was also neglected in this study. Implementation of antibody feedback together with a structural affinity model is computationally challenging as the amount of antibodies produced for every possible conformation needs to be stored to consider epitope-specific antigen masking. The impact of antibody feedback in a murine primary GC response with a short duration is unknown. However, in a secondary GC response and in the case of long-lasting human GCs induced by mRNA vaccines (Turner et al., 2021), antibody feedback could be an important factor modulating the diversity of GC B cells. Indeed, antibody feedback has been shown to alter the generation of memory B cells from GCs by lowering the threshold of B cell activation for GC entry and epitope masking (Inoue et al., 2023). Our simulations show that despite the absence of antibody feedback, subdominant responses can emerge albeit very slowly. This observation suggests that there is a tendency for GC responses to support mild affinity maturation to poorly immunogenic antigens in the presence of highly immunogenic antigens even in conditions with weak antibody feedback. However, the presence of strong antibody feedback might accelerate and enhance the development of subdominant responses (Meyer-Hermann, 2019).

The slow development of immune responses to poorly immunogenic antigens in our simulations suggests that subsequent immunizations might further amplify the subdominant responses, and that timing of immunizations might play an important role in the expansion of responses to low immunogenic antigens. It has indeed been suggested that preserving antibody diversity after the primary immunization is important to increase the chance of raising broadly neutralizing antibodies at subsequent responses (Amitai et al., 2020). Experiments testing iterative immunizations with low immunogenic antigens added either during a primary or secondary immunization at different time-points would inform how the primary response imprints further subdominant responses. Human antibody responses to mRNA vaccines have shown a saturation of antibody levels and limited memory reactivation after the fourth immunization (Pape et al., 2021), which could be a consequence of antibody feedback. However, we showed that GC dynamics are driven by immunodominant responses but do not inform on subdominant concomitant responses. This finding suggests that, despite potential self-inhibition of immunodominant responses by antibody feedback, each subsequent immunization may favor the development of subdominant responses and may still be beneficial despite the lack of observed improved antibody titres.

Increasing the number of antigens was capable of moderately dampening the immunodominant response in our simulations, suggesting that the full antigenic setting should be considered (including potentially bystander antigens) when assessing the relative immunodominance of responses towards two antigens. This observation suggests that antigen-diverse GCs such as GCs present in peyer’s patches (Reboldi & Cyster, 2016) might support the generation of immune responses to antigens with a larger range of immunogenicities in comparison to other lymphoid organs. Although it is well known that mesenteric LN or peyer’s patches provide different (more or less tolerogenic) immune environments (Pezoldt et al., 2018), antigen diversity could be an additional factor explaining the differential response of oral vaccines compared to intradermal vaccines (Stone et al., 2023).

Our study was limited to different domains representing native, non-mutated antigens (i.e., non sequentially related antigens). It is unknown whether combining antigens with differently mutated domains might shift the focus of GCs towards subdominant antigens. When multiple antigens with mutated variants of a highly immunogenic domain are combined, there is a possibility that the GC response to the most immunogenic domain is enhanced due to increased cross-reactivity as observed in our previous study in a single domain setting (Robert et al., 2021). The robustness of immunodominance against variation of antigen amounts observed in the present study, suggests that combining mutated immunogenic domains might therefore have a limited effect in decreasing immunodominance. In contrast, simulations with antigens containing mutated variants of a domain with low immunogenicity would inform whether the induced GC responses would synergize beyond the non-specific effect due to increased antigen valency. Future studies testing these possibilities would be invaluable in the development of strategies to induce the generation of broadly neutralizing antibodies.

## Methods

### Agent-based model of the germinal center

We used *hyphasma*, a previously published agent-based model for the GC response (Meyer-Hermann et al., 2012) that simulates SHM using *Ymir*, a structural representation of antibody-antigen binding (Robert et al., 2021). In the present study, we vary the design of antigens, their amount, and the choice of naive founder cells that enter the GC, and use a fixed mutation rate per amino acid. All other simulation settings are identical to (Robert et al., 2021). A parameter file with the exhaustive list of all used parameters is given in supplementary file 1.

Briefly, cells move on a three-dimensional discrete lattice with nodes of 5 micrometers in each dimension. Every lattice node can only contain one cell. At start, 250 Tfh cells, 200 FDCs with 6 arms (dendrites) following each axis over 40 micrometers are randomly spread in the LZ. Similarly, 300 stromal cells were placed in the DZ. T or B cells are allowed to overlap with the nodes occupied by dendrites of FDCs. The chemokines produced by FDCs and stromal cells are pre-calculated over the GC space and direct the movement of B or T cells toward the DZ and LZ. The simulation uses time-steps of 1e-4 hours. B cells move randomly according to a persistent walk fitted to experimentally observed migration (see (Meyer-Hermann et al., 2012)).

Founder B cells are incorporated randomly at free lattice nodes with an inflow rate of 2 cells per hour with a BCR sequence chosen from a predefined pool (see founder sequences). They proliferate 6 times. Cells divide with a gaussian distribution of cell cycle duration following an average of 7.5 hours per division. During each division, the BCR sequence is randomly mutated (SHM) by uniformly replacing each amino-acid with probability 0.055. Since we simulate sequences of size 9 amino acids, this represents an average mutation probability of 0.5 per division for the full sequence, matching the original simulation settings of *hyphasma*. Of note, multiple point mutations are allowed. Affinity dependent modulation in the mutation probability is not considered in this study, as it is not clear how the affinity of multiple antigens would contribute to this mechanism.

After proliferating in the DZ, B cells migrate to the LZ and are labelled “unselected”. Unselected B cells compete for antigen binding and internalization. The antigens are spread on all nodes covered by FDC dendrites according to the number of antigens considered and the total antigen amount (see distribution of antigens). Every 0.001 hours, the affinity of the BCR of each unselected B cell on a node containing antigen to each of the antigens present in that node is calculated. The B cell captures one unit of the antigen to which the BCR has highest affinity and the probability of antigen capture is proportional to the BCR affinity, provided a minimal affinity of 1e-8. As a comparison, founder cells have BCRs of affinity at least 1e-4. Captured antigen is removed from the node, leading to the consumption of antigen by B cells. An antigen search time of 0.7 hours is allocated for each B cell. Cells that fail to capture any antigen within the antigen search time undergo apoptosis, while the B cells that survive proceed to the Tfh selection phase and initiate contacts with Tfh cells.

Tfh selection is defined with a fixed time-window of 3 hours, with Tfh-B cell polarization time larger than 30 minutes being sufficient for survival within a contact of duration 0.6 hours. At each time-step, it is assumed that Tfh cells only provide surviving signals to the neighbor B cell with the highest amount of internalized antigen. B cells surviving the selection process recycle to the DZ, and are attributed a number of divisions depending on the number of internalized antigens (pMHC-dependent Tfh-induced number of divisions). Of note, internalized antigens are shared asymmetrically between daughter cells in 72% of B cell divisions. At the end of the divisions, B cells retaining the highest amount of antigen become output cells. Output cells differentiate into antibody secreting cells with a rate corresponding to the half-life of 24 hours. Each antibody secreting cell produces 3e-8 moles of antibodies per hour.

### Representation of antibody-antigen affinities

We use a coarse-grained representation of antibodies/BCRs and antigens as lattice proteins (Robert et al., 2021), where amino acids can only occupy integer positions in space, covalently bound amino acids are neighbors, and a lattice position can only contain one amino acid. The experimentally derived interaction potential between amino acids (Miyazawa & Jernigan, 1996) is used to estimate the binding energy between neighboring amino acids of the BCR and the antigen, respectively (Figure 1B). Thanks to the lattice simplification, starting from a BCR sequence and an antigen sequence and structure, every possible binding structure of the BCR around the antigen can be evaluated and given a binding energy (exhaustive docking). The binding structure with optimal energy (including a stabilization factor for energies within the BCR) is defined as the “binding structure” and defines the binding energy of this BCR-antigen pair. Energies are converted into binding affinities using *Aff* = exp((*E* - *E*_max_)/*C*) where *E*_max_ *= -100* and *C = 2*.*8*. Due to high computational costs for calculating binding energies of a new BCR sequence at every SHM event (on average 6.8 million structures to evaluate each time), and the need to use sequences of the same size (longer sequences are energetically favored), we restricted the length of BCR sequences to 9 amino acids, as a model for a substring of the CDRH3 region of the BCR that typically ranges between 10 and 20 amino acids, and which is the determinant region of antibody-antigen binding. This simplification enables the simulation of GC reaction in one CPU within 5 hours per antigen present in the simulation. We have previously shown that this structural representation of antibody-antigen binding models many levels of antibody-antigen binding complexity such as topological hindrance, existence of key mutations, large ranges of affinity, existence of multiple high affinity sequences (Robert et al., 2021), existence of structural domains with higher immunogenicity, and combinatorial amounts of binding modes (Robert et al., 2022).

### Generation of *in silico* antigen structures

We used the discretization method proposed in (Robert et al., 2022) using the *Latfit* software (Mann et al., 2012) to convert the PDB structures of proteins into a lattice representation that minimizes the distance root-mean-square deviations (dRMSD) between the original structure and the lattice structure. We arbitrarily took four antigens from a library of 159 previously discretized antigens (Robert et al., 2022) focusing on simpler antigens with only one chain and less than 160 amino acids to minimize computational costs (ST1: 1OB1 chain C, 159 amino acid; ST2: PDB 1FSK chain A, 95 amino acids; ST3: 1FBI chain X, 122 amino acids; ST3: 1H0D chain C, 129 amino acids). Discretization was performed using Absolut! (*https://github.com/csi-greifflab/Absolut*) with lattice constant 5.25 Å, using the mass center of all atoms in each amino acid (“Fused Centers”).

### Generation of *in silico* antigen sequences and immunogenicity classes

Based on the five scaffold structures ST1 to ST5, we generated 1000 antigens with random amino acid compositions (Figure 1D), and classified the antigens into different immunogenicity classes based on the average binding energy of these antigens to 200 random BCR sequences of 9 amino acids. One antigen per class was selected (Supplementary Table 1). This creates a library of antigens with the same structure but unrelated amino acid sequences (no a priori conservation between them). Simulations are done with antigens of different classes on the same or different scaffold structures.

### Pool of founder sequences

For each antigen separately, within a set of N antigens, a pool of 1000/N naive BCR sequences is generated by randomly sampling sequences of 9 amino acids, and filtering those that have an affinity above 0.0001 to this antigen. A total of 1000 naive sequences are therefore generated, with equal numbers of sequences binding each antigen. Due to possible cross-reactivity, sequences from one antigen pool might recognize other antigens too. During the GC simulation, a BCR sequence is chosen from the pool of sequences for each founder cell.

### Distribution of antigens

A total of 3000 units of antigen are distributed uniformly on every lattice node covered by a FDC dendrite. When multiple antigens are used, their amount is divided equally such that the total amount of antigen is constant.

## Supplementary Figures

**Supplementary Figure 1:**
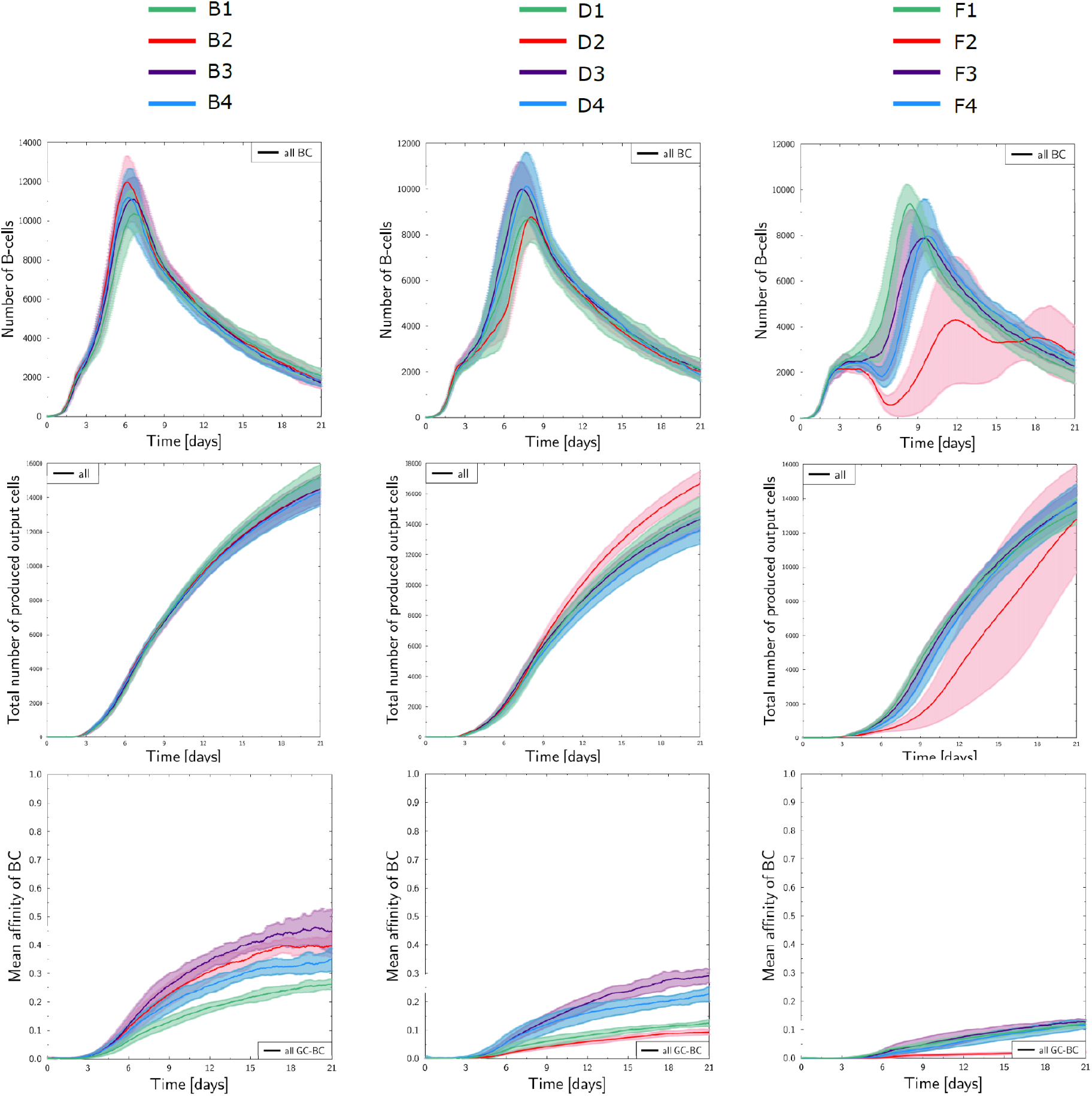
The hierarchy between GC response strength of classes B,D and F is preserved between different scaffold structures. The GC dynamics are shown for an antigen in each immunogenicity class B, D and F, on the four different scaffold structures. Affinities are slightly impacted by the scaffolding structure, but the hierarchy between B, D and F persist within a scaffolding structure.

**Supplementary Figure 2:**
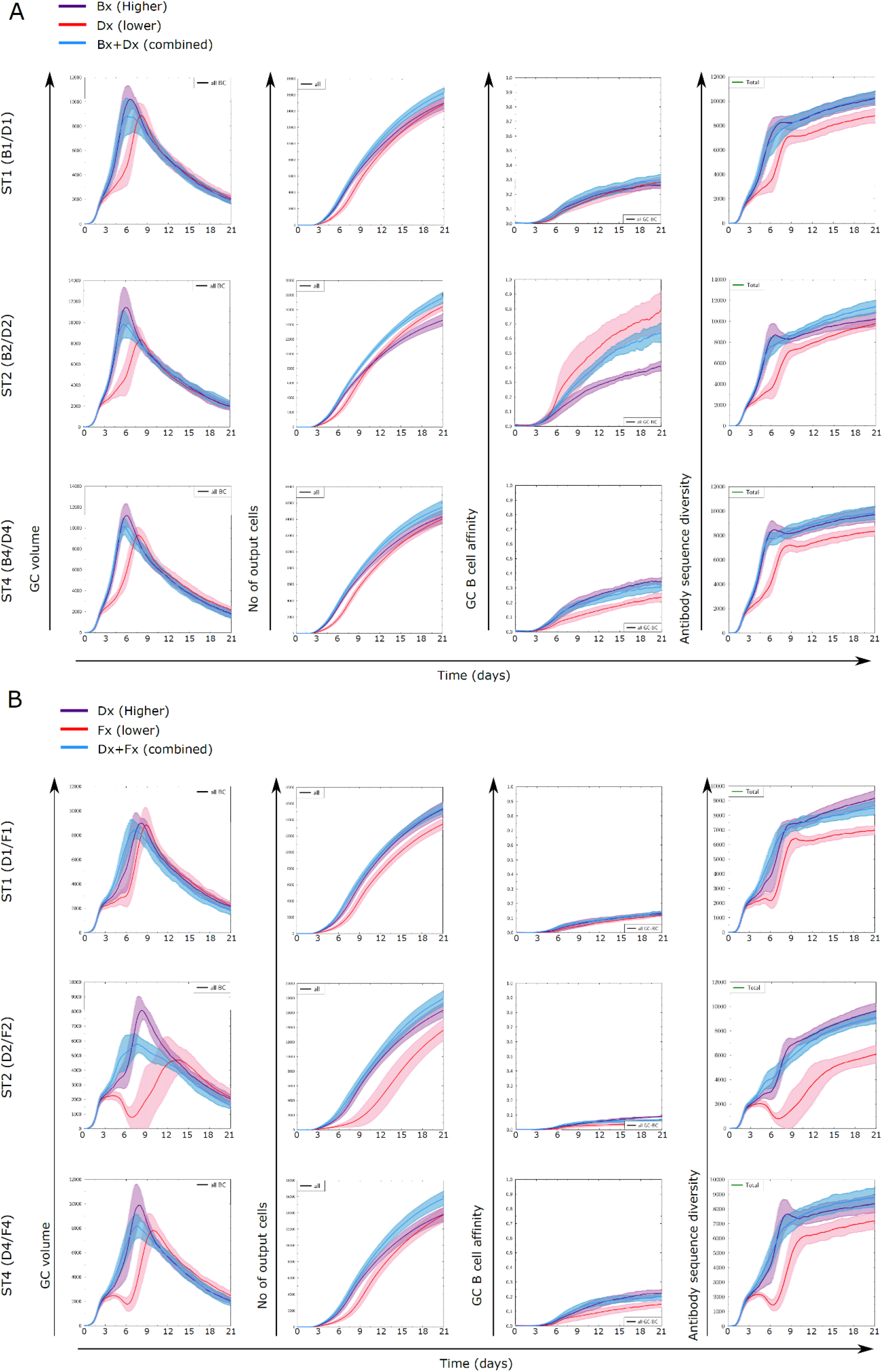
GC dynamics are defined by the antigen of highest immunogenicity; results of Figure 4 are valid on other antigen scaffolds.

**Supplementary Table 1:**
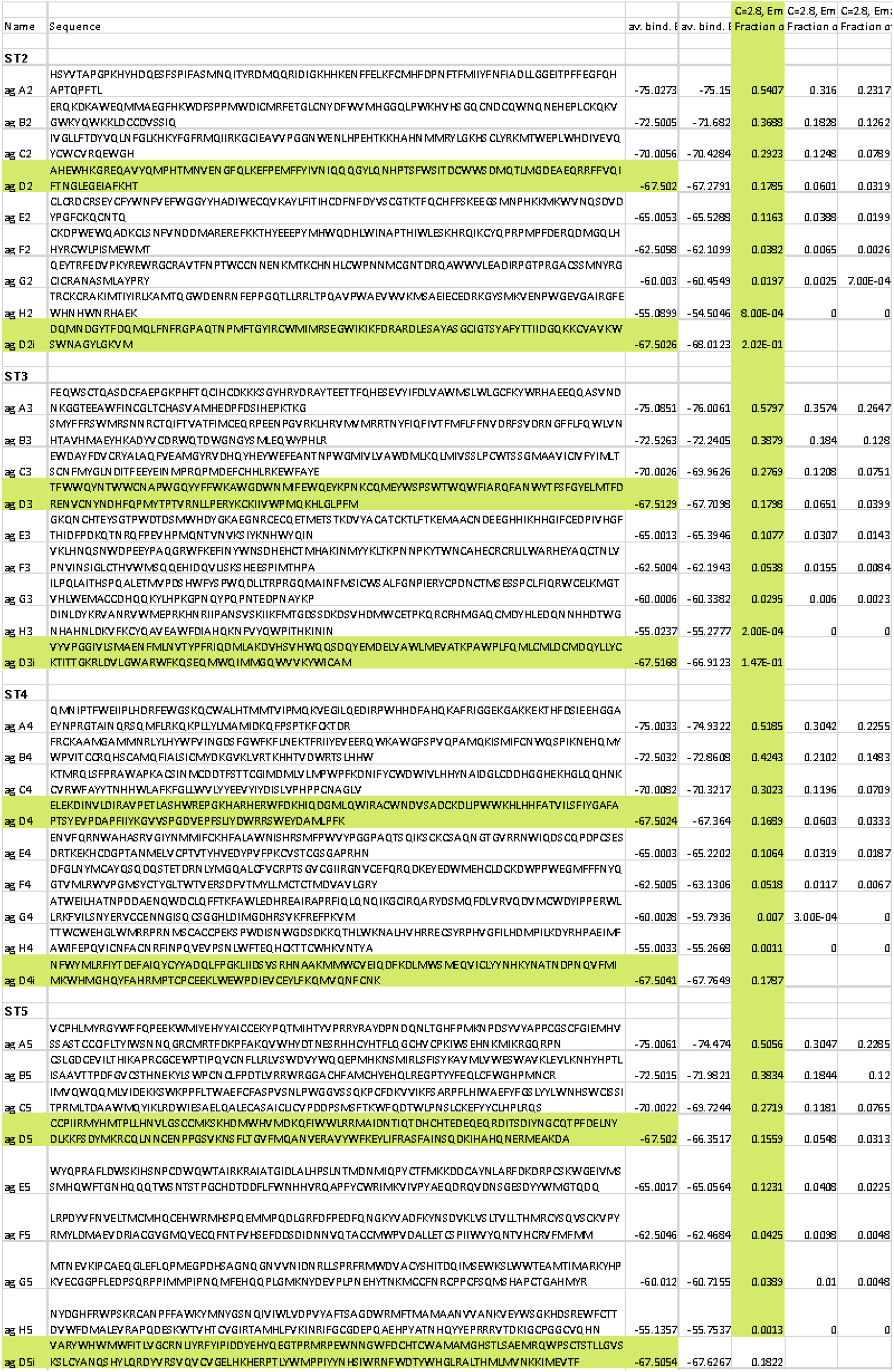
List of antigen sequences used for each scaffold structure. A representative antigen sequence per immunogenicity class (A to H) was taken, the number denotes the scaffolding structure. Antigens Di are generated for simulations with two antigens of class D on the same scaffold, preserving the average binding energy.

**Supplementary File 1: A reference parameter file**

## References

Abbott, R. K., Lee, J. H., Menis, S., Skog, P., Rossi, M., Ota, T., Kulp, D. W., Bhullar, D., Kalyuzhniy, O., Havenar-Daughton, C., Schief, W. R., Nemazee, D., & Crotty, S. (2018). Precursor Frequency and Affinity Determine B Cell Competitive Fitness in Germinal Centers, Tested with Germline-Targeting HIV Vaccine Immunogens. /u>Immunity, 48(1). https://doi.org/10.1016/j.immuni.2017.11.023

Allen, C. D., Okada, T., & Cyster, J. G. (2007). Germinal-center organization and cellular dynamics. Immunity, 27(2). https://doi.org/10.1016/j.immuni.2007.07.009

Amitai, A., Sangesland, M., Barnes, R. M., Rohrer, D., Lonberg, N., Lingwood, D., & Chakraborty, A. K. (2020). Defining and Manipulating B Cell Immunodominance Hierarchies to Elicit Broadly Neutralizing Antibody Responses against Influenza Virus. Cell Systems, 11(6). https://doi.org/10.1016/j.cels.2020.09.005

Baumjohann, D., Preite, S., Reboldi, A., Ronchi, F., Ansel, K. M., Lanzavecchia, A., & Sallusto, F. (2013). Persistent antigen and germinal center B cells sustain T follicular helper cell responses and phenotype. Immunity, 38(3). https://doi.org/10.1016/j.immuni.2012.11.020

Belongia, E. A., Simpson King, J. P., Sundaram, M. E., Kelley, N. S., Osterholm, M. T., & McLean, H. Q. (2016). Variable influenza vaccine effectiveness by subtype: a systematic review and meta-analysis of test-negative design studies. The Lancet Infectious Diseases, 16(8). https://doi.org/10.1016/S1473-3099(16)00129-8

Buchauer, L., & Wardemann, H. (2019). Calculating germinal centre reactions. Current Opinion in Systems Biology, 18, 1–8.

Burnett, D. L., Langley, D. B., Schofield, P., Hermes, J. R., Chan, T. D., Jackson, J., Bourne, K., Reed, J. H., Patterson, K., Porebski, B. T., Brink, R., Christ, D., & Goodnow, C. C. (2018). Germinal center antibody mutation trajectories are determined by rapid self/foreign discrimination. Science, 360(6385). https://doi.org/10.1126/science.aao3859

Carter, D. M., Darby, C. A., Lefoley, B. C., Crevar, C. J., Alefantis, T., Oomen, R., Anderson, S. F., Strugnell, T., Cortés-Garcia, G., Vogel, T. U., Parrington, M., Kleanthous, H., & Ross, T. M. (2016). Design and Characterization of a Computationally Optimized Broadly Reactive Hemagglutinin Vaccine for H1N1 Influenza Viruses. Journal of Virology, 90(9). https://doi.org/10.1128/JVI.03152-15

Chai, N., Swem, L. R., Reichelt, M., Chen-Harris, H., Luis, E., Park, S., Fouts, A., Lupardus, P., Wu, T. D., Li, O., McBride, J., Lawrence, M., Xu, M., & Tan, M.-W. (2016). Two Escape Mechanisms of Influenza A Virus to a Broadly Neutralizing Stalk-Binding Antibody. PLoS Pathogens, 12(6), e1005702.

Childs, L. M., Baskerville, E. B., & Cobey, S. (2015). Trade-offs in antibody repertoires to complex antigens. Philosophical Transactions of the Royal Society of London. Series B, Biological Sciences, 370(1676). https://doi.org/10.1098/rstb.2014.0245

Doud, M. B., Lee, J. M., & Bloom, J. D. (2018). How single mutations affect viral escape from broad and narrow antibodies to H1 influenza hemagglutinin. Nature Communications, 9(1), 1–12.

Escolano, A., Steichen, J. M., Dosenovic, P., Kulp, D. W., Golijanin, J., Sok, D., Freund, N. T., Gitlin, A. D., Oliveira, T., Araki, T., Lowe, S., Chen, S. T., Heinemann, J., Yao, K. H., Georgeson, E., Saye-Francisco, K. L., Gazumyan, A., Adachi, Y., Kubitz, M.,… Nussenzweig, M. C. (2016). Sequential Immunization Elicits Broadly Neutralizing Anti-HIV-1 Antibodies in Ig Knockin Mice. Cell, 166(6). https://doi.org/10.1016/j.cell.2016.07.030

Finney, J., Yeh, C. H., Kelsoe, G., & Kuraoka, M. (2018). Germinal center responses to complex antigens. Immunological Reviews, 284(1). https://doi.org/10.1111/imr.12661

Francis, T. (1960). On the doctrine of original antigenic sin. Proc. Am. Philos. Soc., 104, 572–578.

Gao, F., Bonsignori, M., Liao, H.-X., Kumar, A., Xia, S.-M., Lu, X., Cai, F., Hwang, K.-K., Song, H., Zhou, T., Lynch, R. M., Munir Alam, S., Anthony Moody, M., Ferrari, G., Berrong, M., Kelsoe, G., Shaw, G. M., Hahn, B. H., Montefiori, D. C.,… Haynes, B. F. (2014). Cooperation of B Cell Lineages in Induction of HIV-1-Broadly Neutralizing Antibodies. Cell, 158(3), 481–491.

Garg, A. K., Mitra, T., Schips, M., Bandyopadhyay, A., & Meyer-Hermann, M. (2022). Amount of antigen, T follicular helper cells and quality of seeder cells shape the diversity of germinal center B cells. In bioRxiv (p. 2022.10.26.513835). https://doi.org/10.1101/2022.10.26.513835

Garg, A. K., Mitra, T., Schips, M., Bandyopadhyay, A., & Meyer-Hermann, M. (2023). Amount of antigen, T follicular helper cells and affinity of founder cells shape the diversity of germinal center B cells: A computational study. Frontiers in Immunology, 14. https://doi.org/10.3389/fimmu.2023.1080853

Gitlin, A. D., Mayer, C. T., Oliveira, T. Y., Shulman, Z., Jones, M. J. K., Koren, A., & Nussenzweig, M. C. (2015). T cell help controls the speed of the cell cycle in germinal center B cells. Science, 349(6248), 643.

Haynes, B. F., Wiehe, K., Borrrow, P., Saunders, K. O., Korber, B., Wagh, K., McMichael, A. J., Kelsoe, G., Hahn, B. H., Alt, F., & Shaw, G. M. (2022). Strategies for HIV-1 vaccines that induce broadly neutralizing antibodies. Nature Reviews. Immunology, 1–17.

Ilinskaya, A. N., & Dobrovolskaia, M. A. (2016). Understanding the immunogenicity and antigenicity of nanomaterials: Past, present and future. Toxicology and Applied Pharmacology, 299, 70.

Inoue, T., Shinnakasu, R., Kawai, C., Yamamoto, H., Sakakibara, S., Ono, C., Itoh, Y., Terooatea, T., Yamashita, K., Okamoto, T., Hashii, N., Ishii-Watabe, A., Butler, N. S., Matsuura, Y., Matsumoto, H., Otsuka, S., Hiraoka, K., Teshima, T., Murakami, M., & Kurosaki, T. (2023). Antibody feedback contributes to facilitating the development of Omicron-reactive memory B cells in SARS-CoV-2 mRNA vaccinees. The Journal of Experimental Medicine, 220(2). https://doi.org/10.1084/jem.20221786

Kuraoka, M., Schmidt, A. G., Nojima, T., Feng, F., Watanabe, A., Kitamura, D., Harrison, S. C., Kepler, T. B., & Kelsoe, G. (2016). Complex Antigens Drive Permissive Clonal Selection in Germinal Centers. Immunity, 44(3). https://doi.org/10.1016/j.immuni.2016.02.010

Liao, H. X., Lynch, R., Zhou, T., Gao, F., Alam, S. M., Boyd, S. D., Fire, A. Z., Roskin, K. M., Schramm, C. A., Zhang, Z., Zhu, J., Shapiro, L., Mullikin, J. C., Gnanakaran, S., Hraber, P., Wiehe, K., Kelsoe, G., Yang, G., Xia, S. M.,… Haynes, B. F. (2013). Co-evolution of a broadly neutralizing HIV-1 antibody and founder virus. Nature, 496(7446). https://doi.org/10.1038/nature12053

Linterman, M., Silva-Cayetano, A., Fra-Bido, S., Innocentin, S., Lee, J. L., Webb, L., Foster, W., Burton, A., Bignon, A., Vanderleyden, I., Carr, E., Hill, D., Denton, A., Robert, P., Meyer-Hermann, M., Alterauge, D., Baumjohann, D., Lemos, J., Espeli, M., & Balabanian, K. (2023). Spatial dysregulation of T follicular helper cells impairs vaccine responses in ageing. Nature Immunology. https://doi.org/10.21203/rs.3.rs-1733421/v1

Mann, M., Saunders, R., Smith, C., Backofen, R., & Deane, C. M. (2012). Producing high-accuracy lattice models from protein atomic coordinates including side chains. Advances in Bioinformatics, 2012. https://doi.org/10.1155/2012/148045

McNamara, H. A., Idris, A. H., Sutton, H. J., Vistein, R., Flynn, B. J., Cai, Y., Wiehe, K., Lyke, K. E., Chatterjee, D., Kc, N., Chakravarty, S., Bk, L. S., Hoffman, S. L., Bonsignori, M., Seder, R. A., & Cockburn, I. A. (2020). Antibody Feedback Limits the Expansion of B Cell Responses to Malaria Vaccination but Drives Diversification of the Humoral Response. Cell Host & Microbe, 28(4). https://doi.org/10.1016/j.chom.2020.07.001

Mesin, L., Schiepers, A., Ersching, J., Barbulescu, A., Cavazzoni, C. B., Angelini, A., Okada, T., Kurosaki, T., & Victora, G. D. (2020). Restricted Clonality and Limited Germinal Center Reentry Characterize Memory B Cell Reactivation by Boosting. Cell, 180(1). https://doi.org/10.1016/j.cell.2019.11.032

Meyer-Hermann, M. (2019). Injection of Antibodies against Immunodominant Epitopes Tunes Germinal Centers to Generate Broadly Neutralizing Antibodies. Cell Reports, 29(5), 1066–1073.e5.

Meyer-Hermann, M. (2021). A molecular theory of germinal center B cell selection and division. Cell Reports, 36(8). https://doi.org/10.1016/j.celrep.2021.109552

Meyer-Hermann, M., Mohr, E., Pelletier, N., Zhang, Y., Victora, G. D., & Toellner, K. M. (2012). A theory of germinal center B cell selection, division, and exit. Cell Reports, 2(1). https://doi.org/10.1016/j.celrep.2012.05.010

Miyazawa, S., & Jernigan, R. L. (1996). Residue-residue potentials with a favorable contact pair term and an unfavorable high packing density term, for simulation and threading. Journal of Molecular Biology, 256(3). https://doi.org/10.1006/jmbi.1996.0114

Newman, J., Thakur, N., Peacock, T. P., Bialy, D., Elrefaey, A. M. E., Bogaardt, C., Horton, D. L., Ho, S., Kankeyan, T., Carr, C., Hoschler, K., Barclay, W. S., Amirthalingam, G., Brown, K. E., Charleston, B., & Bailey, D. (2022). Neutralizing antibody activity against 21 SARS-CoV-2 variants in older adults vaccinated with BNT162b2. Nature Microbiology, 7(8), 1180–1188.

Pape, K. A., Dileepan, T., Kabage, A. J., Kozysa, D., Batres, R., Evert, C., Matson, M., Lopez, S., Krueger, P. D., Graiziger, C., Vaughn, B. P., Shmidt, E., Rhein, J., Schacker, T. W., Khoruts, A., & Jenkins, M. K. (2021). High-affinity memory B cells induced by SARS-CoV-2 infection produce more plasmablasts and atypical memory B cells than those primed by mRNA vaccines. Cell Reports, 37(2). https://doi.org/10.1016/j.celrep.2021.109823

Pezoldt, J., Pasztoi, M., Zou, M., Wiechers, C., Beckstette, M., Thierry, G. R., Vafadarnejad, E., Floess, S., Arampatzi, P., Buettner, M., Schweer, J., Fleissner, D., Vital, M., Pieper, D. H., Basic, M., Dersch, P., Strowig, T., Hornef, M., Bleich, A.,… Huehn, J. (2018). Neonatally imprinted stromal cell subsets induce tolerogenic dendritic cells in mesenteric lymph nodes. Nature Communications, 9(1), 1–14.

Qiu, Y., Stegalkina, S., Zhang, J., Boudanova, E., Park, A., Zhou, Y., Prabakaran, P., Pougatcheva, S., Ustyugova, I. V., Vogel, T. U., Mundle, S. T., Oomen, R., Delagrave, S., Ross, T. M., Kleanthous, H., & Qiu, H. (2020). Mapping of a Novel H3-Specific Broadly Neutralizing Monoclonal Antibody Targeting the Hemagglutinin Globular Head Isolated from an Elite Influenza Virus-Immunized Donor Exhibiting Serological Breadth. Journal of Virology, 94(6). https://doi.org/10.1128/JVI.01035-19

Reboldi, A., & Cyster, J. G. (2016). Peyer’s patches: Organizing B cell responses at the intestinal frontier. Immunological Reviews, 271(1), 230.

Robert, P. A., Akbar, R., Frank, R., Pavlović, M., Widrich, M., Snapkov, I., Slabodkin, A., Chernigovskaya, M., Scheffer, L., Smorodina, E., Rawat, P., Mehta, B. B., Vu, M. H., Mathisen, I. F., Prósz, A., Abram, K., Olar, A., Miho, E., Haug, D. T. T.,… Greiff, V. (2022). Unconstrained generation of synthetic antibody–antigen structures to guide machine learning methodology for antibody specificity prediction. Nature Computational Science, 2(12), 845–865.

Robert, P. A., Arulraj, T., & Meyer-Hermann, M. (2021). Ymir: A 3D structural affinity model for multi-epitope vaccine simulations. iScience, 24(9), 102979.

Robert, P. A., Marschall, A. L., & Meyer-Hermann, M. (2018). Induction of broadly neutralizing antibodies in Germinal Centre simulations. Current Opinion in Biotechnology, 51. https://doi.org/10.1016/j.copbio.2018.01.006

Saunders, K. O., Nicely, N. I., Wiehe, K., Bonsignori, M., Meyerhoff, R. R., Parks, R., Walkowicz, W. E., Aussedat, B., Wu, N. R., Cai, F., Vohra, Y., Park, P. K., Eaton, A., Go, E. P., Sutherland, L. L., Scearce, R. M., Barouch, D. H., Zhang, R., Von Holle, T.,… Haynes, B. F. (2017). Vaccine Elicitation of High Mannose-Dependent Neutralizing Antibodies against the V3-Glycan Broadly Neutralizing Epitope in Nonhuman Primates. Cell Reports, 18(9). https://doi.org/10.1016/j.celrep.2017.02.003

Sheng, J., & Wang, S. (2021). Coevolutionary transitions emerging from flexible molecular recognition and eco-evolutionary feedback. iScience, 24(8), 102861.

Shlomchik, M. J., & Weisel, F. (2012). Germinal center selection and the development of memory B and plasma cells. Immunological Reviews, 247(1). https://doi.org/10.1111/j.1600-065X.2012.01124.x

Silver, J., Zuo, T., Chaudhary, N., Kumari, R., Tong, P., Giguere, S., Granato, A., Donthula, R., Devereaux, C., & Wesemann, D. R. (2018). Stochasticity enables BCR-independent germinal center initiation and antibody affinity maturation. The Journal of Experimental Medicine, 215(1). https://doi.org/10.1084/jem.20171022

Sokal, A., Barba-Spaeth, G., Hunault, L., Fernández, I., Broketa, M., Meola, A., Fourati, S., Azzaoui, I., Vandenberghe, A., Lagouge-Roussey, P., Broutin, M., Roeser, A., Bouvier-Alias, M., Crickx, E., Languille, L., Fournier, M., Michel, M., Godeau, B., Gallien, S.,… Chappert, P. (2023). Omicron BA.1 breakthrough infection drives long-term remodeling of the memory B cell repertoire in vaccinated individuals. In bioRxiv (p. 2023.01.27.525575). https://doi.org/10.1101/2023.01.27.525575

Stebegg, M., Kumar, S. D., Silva-Cayetano, A., Fonseca, V. R., Linterman, M. A., & Graca, L. (2018). Regulation of the Germinal Center Response. Frontiers in Immunology, 9. https://doi.org/10.3389/fimmu.2018.02469

Stone, A. E., Rambaran, S., Trinh, I. V., Estrada, M., Jarand, C. W., Williams, B. S., Murrell, A. E., Huerter, C. M., Bai, W., Palani, S., Nakanishi, Y., Laird, R. M., Poly, F. M., Reed, W. F., White, J. A., & Norton, E. B. (2023). Route and antigen shape immunity to dmLT-adjuvanted vaccines to a greater extent than biochemical stress or formulation excipients. Vaccine, 41(9). https://doi.org/10.1016/j.vaccine.2023.01.033

Tas, J. M., Mesin, L., Pasqual, G., Targ, S., Jacobsen, J. T., Mano, Y. M., Chen, C. S., Weill, J. C., Reynaud, C. A., Browne, E. P., Meyer-Hermann, M., & Victora, G. D. (2016). Visualizing antibody affinity maturation in germinal centers. Science, 351(6277). https://doi.org/10.1126/science.aad3439

Tian, M., Cheng, C., Chen, X., Duan, H., Cheng, H. L., Dao, M., Sheng, Z., Kimble, M., Wang, L., Lin, S., Schmidt, S. D., Du, Z., Joyce, M. G., Chen, Y., DeKosky, B. J., Chen, Y., Normandin, E., Cantor, E., Chen, R. E.,… Alt, F. W. (2016). Induction of HIV Neutralizing Antibody Lineages in Mice with Diverse Precursor Repertoires. Cell, 166(6). https://doi.org/10.1016/j.cell.2016.07.029

Turner, J. S., O’Halloran, J. A., Kalaidina, E., Kim, W., Schmitz, A. J., Zhou, J. Q., Lei, T., Thapa, M., Chen, R. E., Case, J. B., Amanat, F., Rauseo, A. M., Haile, A., Xie, X., Klebert, M. K., Suessen, T., Middleton, W. D., Shi, P.-Y., Krammer, F.,… Ellebedy, A. H. (2021). SARS-CoV-2 mRNA vaccines induce persistent human germinal centre responses. Nature, 596(7870), 109–113.

Victora, G. D., & Nussenzweig, M. C. (2022). Germinal Centers. Annual Review of Immunology, 40. https://doi.org/10.1146/annurev-immunol-120419-022408

Wang, S. (2017). Optimal Sequential Immunization Can Focus Antibody Responses against Diversity Loss and Distraction. PLoS Computational Biology, 13(1), e1005336.

Wang, S., Mata-Fink, J., Kriegsman, B., Hanson, M., Irvine, D. J., Eisen, H. N., Burton, D. R., Wittrup, K. D., Kardar, M., & Chakraborty, A. K. (2015). Manipulating the selection forces during affinity maturation to generate cross-reactive HIV antibodies. Cell, 160(4). https://doi.org/10.1016/j.cell.2015.01.027

Wanzeck, K., Boyd, K. L., & McCullers, J. A. (2011). Glycan Shielding of the Influenza Virus Hemagglutinin Contributes to Immunopathology in Mice. American Journal of Respiratory and Critical Care Medicine, 183(6), 767.

Yan, L., & Wang, S. (2020). Shaping Polyclonal Responses via Antigen-Mediated Antibody Interference. iScience, 23(10), 101568.

Zhang, Y., Garcia-Ibanez, L., & Toellner, K. M. (2016). Regulation of germinal center B-cell differentiation. Immunological Reviews, 270(1). https://doi.org/10.1111/imr.12396

Zhang, Y., Meyer-Hermann, M., George, L. A., Figge, M. T., Khan, M., Goodall, M., Young, S. P., Reynolds, A., Falciani, F., Waisman, A., Notley, C. A., Ehrenstein, M. R., Kosco-Vilbois, M., & Toellner, K.-M. (2013). Germinal center B cells govern their own fate via antibody feedback. The Journal of Experimental Medicine, 210(3), 457–464.

